# Overexpression of flavodiiron protein Flv3 in engineered *Synechocystis* stimulates sucrose production and growth by altering cellular redox balance through enhanced sulfur metabolism

**DOI:** 10.64898/2026.06.23.733971

**Authors:** R. Ndeh, D. Muth-Pawlak, E. Moser, A. Tiwari, E-M. Aro, P. Kallio

## Abstract

Biotechnological applications of oxygenic photosynthetic organisms depend on conversion of light energy into chemical energy through photosystems (PS). This energy can then be used to drive engineered metabolic pathways that are designed as strong electron sinks. For optimal performance, the engineered host metabolism must also be balanced with the native photoprotective electron transfer network. This includes the energy-consuming function of flavodiiron (Flv) proteins, which are universal to cyanobacteria and all other oxygenic photosynthetic organisms except angiosperms. In the cyanobacterium *Synechocystis* sp. PCC 6803, four different Flv proteins have been shown to function in a Mehler-like reaction within two heterodimeric forms (Flv1/Flv3 and Flv2/Flv4), donating electrons to O_2_ without generating oxidative stress. Previously, deleting Flv3 in the *Synechocystis* sucrose-producing (S02) strain was shown to cause drastic metabolic changes in S02Δ*flv3,* shifting it from photoautotrophic to mixotrophic growth (Muth-Pawlak, et al., 2024). In this study, we took an opposite approach by complementing S02 with Flv3 overexpression at different levels using RBS tuning. Interestingly, this resulted in S02oeFlv3 strains with significantly increased overall photosynthetic activity and sucrose production, enhanced cell growth, and storage compound accumulation. However, these outcomes are shown not to be due to conventional O_2_ photoreduction activity catalysed by Flv1/Flv3. Instead, we postulate that the observed changes are linked to the previously unidentified function of homomeric Flv3/Flv3 and the strongly increased sulphate redox metabolism. Based on extensive proteomic and metabolite analyses, we hypothesise that the Flv3 homooligomer uses sulfate metabolites directly or indirectly as the final electron acceptor instead of O_2_. This would also explain the upregulation of sulfate-related enzymes, as well as SQR, which passes the electrons back to the PQ pool in the Flv3 overexpression strain.

## INTRODUCTION

*Synechocystis* sp. PCC 6803 (hereafter *Synechocystis*) is a model photosynthetic prokaryote often used as a chassis for cyanobacteria cell factories. It uses H_2_O as an electron source to convert solar energy into chemical energy. ATP (an energy carrier) and NADPH (a reducing agent) are generated via photosynthetic light reactions and are primarily used for carbon assimilation in the Calvin-Benson-Bassham (CBB) cycle and other metabolic pathways. A significant proportion of the converted energy is also utilised by regulatory systems that protect the thylakoid-associated photosynthetic apparatus from the harmful effects of light energy when there is an imbalance between light reactions and cellular metabolism.

Flavodiiron proteins (Flvs) are an important regulatory system of photosynthetic light reactions in cyanobacteria. They direct the surplus of electrons generated in the photosynthetic electron transfer chain towards molecular oxygen, thereby protecting the photosynthetic components from overreduction and photodamage (Allahverdiyeva, et al., 2013) (Helman, et al., 2003) (Zhang, et al., 2009) (Zhang, et al., 2012). In *Synechocystis,* four genes encode the Flv1, Flv2, Flv3 and Flv4 proteins, which function as Flv1/Flv3 and Flv2/Flv4 heterodimers. The expression levels of Flv1-4 proteins depend on environmental conditions: Flv1 and Flv3 are always expressed in the cells, whereas Flv2 and Flv4 are specifically induced when the cells experience low ambient carbon conditions. Notably, Flv2 and Flv4 are encoded in the same operon in *Synechocystis*, unlike Flv3 and Flv1, which are encoded by separate genes (sll0550 and sll0219, respectively) under independent promotors. This independent control may be relevant to the physiological function of the proteins, reflecting the importance of expressing the interacting Flv1 and Flv3 proteins at different levels. Indeed, it has been hypothesized that the Flv3 proteins in particular form different homooligomers but their function remains unclear (Mustila, et al., 2016).

Electrons for reduction of the Flv1/Flv3 heterodimers are derived from PSI, apparently via ferredoxin (Fd), and reduced Flv1/Flv3 oligomers then donate electrons to molecular oxygen and produce water, without generating reactive oxygen species (ROS) (Helman, et al., 2003) (Sétif, et al., 2020). This process, known as the Mehler-like reaction (Allahverdiyeva, et al., 2011) safeguards PSI from photodamage (Allahverdiyeva, et al., 2013) (Mustila, et al., 2016) (Santana Sanchez, et al., 2019), particularly under fluctuating light conditions. Even though the structural and functional properties of Flv proteins have been extensively studied during the past 10 to 20 years, it is highly conceivable that they also carry functions other than the Mehler-like reaction, possibly depending on oligomerization properties, which still remain to be discovered.

Flvs are an important target for biotechnological applications since they redirect the electrons from PSI via Fd to reduce oxygen in an apparently futile cycle and thereby, from the production efficiency viewpoint, waste the energy already harnessed by cyanobacteria. The efficiency of photosynthesis relies on the balance between ATP and NADPH generation and CO_2_ fixation rates as well as on further steps of carbon, nitrogen and sulfur assimilation into final products. Engineering of the Flv functions in cyanobacteria cell factory may enable to direct carbon flow towards desirable compounds (sink) and improve photosynthetic activity by alleviation of reductive pressure from PSI (Hasunuma, et al., 2014) (Santos-Merino, et al., 2021) (Abramson, et al., 2016). The deletion of the *flv3* gene from engineered sucrose producing and excreting *Synechocystis* strain resulted in high rate of sucrose production in an early phase of cultivation, but soon switched to mixotrophic growth mode and sucrose consumption (Thiel, et al., 2019). The clear signs of deteriorated photosynthesis and increase in carbohydrate catabolic reactions proved that the introduced sucrose pathway did not function properly in the absence of Flv3 (Muth-Pawlak, et al., 2024). These experiments proved that inactivation of the *flv3* gene sparked unprecedented metabolic rearrangements leading towards new equilibrium between production of reducing power and its use in *Synechocystis* metabolism.

Here, we took an opposite approach and complemented the *Synechocystis* sucrose production strain (S02) (Thiel, et al., 2019) with overexpression of the Flv3 protein (S02oeFlv3). We investigated the influence of increasing amount of Flv3 on phenotypic and molecular features of engineered *Synechosystis* strain. Comparison of the highest Flv3 overproduction strain with the control sucrose-producing S02 reference showed the sucrose pathway to function as an efficient electron sink. This was accompanied by drastic proteome- and metabolic-level changes, as well as an upregulation of a putative Flv3 homooligomer-mediated electron transfer related to enhanced sulfur metabolism.

## MATERIALS AND METHODS

### Reagents and enzymes

The enzymes utilized in this work were obtained from New England BioLabs (US) or Thermo Fisher Scientific (US), and commercial kits for plasmid extraction and gel isolation from Qiagen (DE). Oligonucleotides were ordered from Eurofins MWG Operon (DE), while larger gene fragments were acquired from GenScript (US). All chemicals used in this study were purchased from Sigma-Aldrich (US), unless stated otherwise.

### *E. coli* strains and culture conditions

*E. coli* strain DH5α was utilized for all DNA cloning procedures and plasmid amplification. The cultures were grown at 37°C in Lysogeny Broth (LB) liquid medium (150–200 rpm agitation), or on 1.5% agar LB plates. For selection the media contained either 100 µg/ml ampicillin (pUC57, pNiv), 50 µg/ml spectinomycin with 20 µg/ml streptomycin (pDF empty) or 50 µg/ml spectinomycin and 34 µg/ml chloramphenicol (pDF with insert). *E. coli* transformants were preserved at –80°C as 10% glycerol stocks.

### Cyanobacterial strains and culture conditions

*Synechocystis sp*. PCC 6803 Δ*ggpS* (Thiel, et al., 2019) was used as background for all the strains generated in this study (**Table 1**). The strains were cultivated in liquid BG11 medium buffered with 20 mM TES-KOH (pH 8.0) with supplemented spectinomycin (Sp; 20 µg/ml) and chloramphenicol (Cm; 8 µg/ml). The liquid cultures were carried out in 250 ml Erlenmeyer flasks containing 100 ml of BG11 medium, incubated at 30 °C with shaking at ∼120 rpm, under continuous lighting of 50–200 µmol photons m^−2^ s^−1^ and 1% CO_2_ atmosphere. A custom-engineered LED lighting system (Kakko, et al., Preprint) was employed to maintain consistent light conditions across all parallel cultures. Pre-cultures were initially grown under continuous illumination of 50 µmol photons m^−2^ s^-1^ until OD_750_∼2. The cultures were then diluted in fresh medium to OD_750_∼0.4, partitioned into four parallel replicates with supplemented 400 mM NaCl under 200 µmol photons m^−2^ s^-1^ for 24 h. Following this acclimation phase, the main cultures were adjusted to OD_750_∼0.5 by gentle pelleting and resuspension in 100 ml of fresh BG-11 medium containing 400 mM NaCl and 1 mM IPTG. Cultures were then incubated under 200 µmol photons m^−2^ s^-1^ for a duration of 12 days.

**Table 1.**
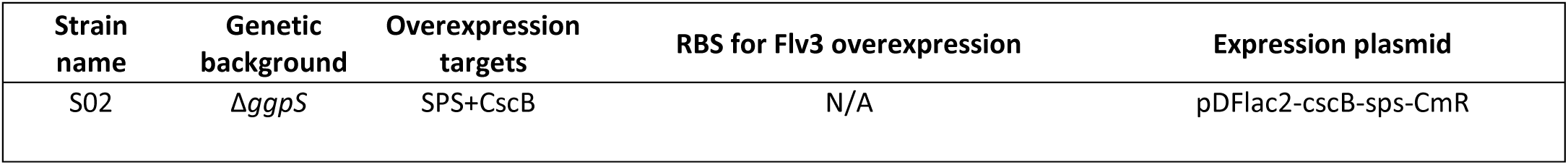
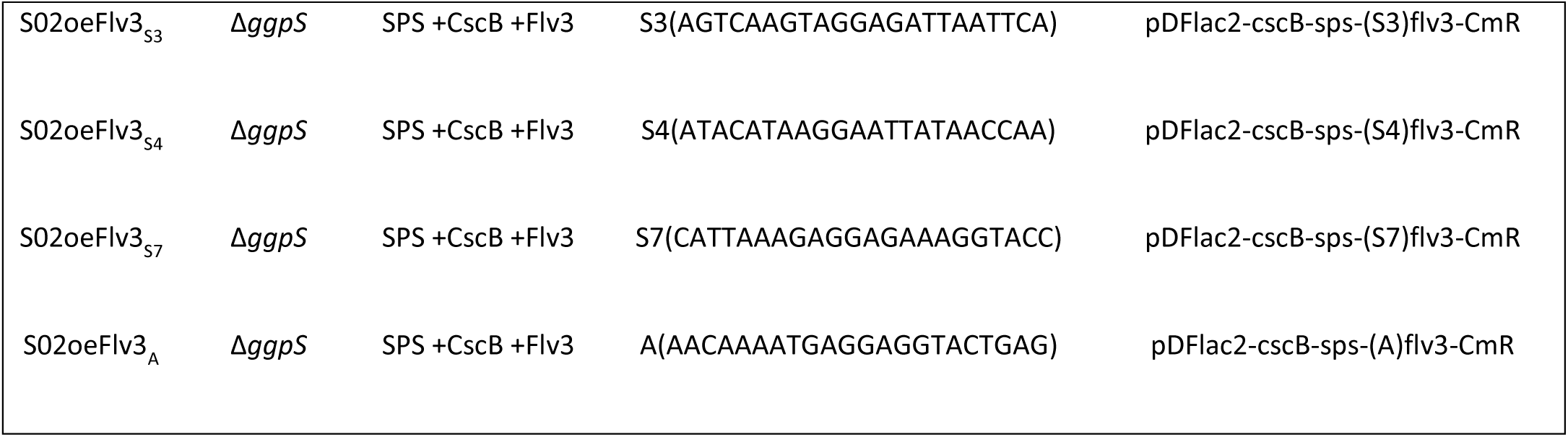
Description of the engineered sucrose-producing *Synechocystis* strains compared in the study. The previously engineered control strain S02 (Thiel, et al., 2019) used as reference overexpresses the sucrose phosphate synthase (SPS) and sucrose permease (CscB; *E. coli*) in the glucosylglycerol-phosphate deletion (Δ*ggps*) background. The derived Flv3 overexpression strains generated in this work additionally overexpress the endogenous flavodiiron protein 3 (Flv3) under the translational control of four alternative RBS elements S3, S4, S7 and A (Thiel, et al., 2018).

### Monitoring cell growth

The growth of the strains was monitored spectrophotometrically by measuring optical density at 600 nm (OD_600n_; *E. coli*) or 750 nm (OD_750_; *Synechocystis*) using GENESYS 10S UV–Vis spectrophotometer (Thermo Fisher Scientific, US). The OD values were calculated from culture dilutions at the linear range of the spectrophotometer (∼0.1-0.5).

### Assembly of Flv3 overexpression constructs

The expression constructs were assembled using a modular subcloning procedure (Thiel, et al., 2018) (Zelcbuch, et al., 2013) where each target gene is first fused with a selected translational control element (RBS), combined into polycistronic operons, and finally transferred into the expression plasmid backbone. Following this strategy, the *Synechocystis flv3* (*sll0550*) coding sequence was ordered as a synthetic fragment in pUC57 backbone (Genscript) with NsiI/XhoI overhangs and subcloned into four alternative pNiv assembly vectors carrying different RBSs (S3, S4, S7, and A) (Thiel, et al., 2018). The resulting RBS-*flv3* fragments (SpeI/SalI) were then fused with the existing sucrose operon pNiv-(S3)cscB-(S3)sps-CmR (NheI/XhoI) (Thiel, et al., 2019) to generate pNiv-(S3)cscB-(S3)sps-(S3/S4/S7/A)flv3-CmR. In the last step, the complete inserts carrying the three target genes were subcloned into the pDF-lac2 expression plasmid backbone (SpeI/SalI) to produce the four final constructs listed in **Table 1**.

### *Synechocystis* transformation and recombinant strain verification

The *Synechocystis* Δ*ggpS* host strain was transformed with the generated expression constructs (**Table 1**) via natural transformation at the logarithmic growth phase (OD_750_ =0.8) by resuspending 5 ml of freshly grown cell culture (OD_750_∼0.8) in 1 ml BG11 together with ∼20μg plasmid DNA extracted from *E. coli*. After o/n incubation in gentle shaking in dark, the cells were transferred to BG-11 agar plates containing 3 μg ml^−1^ chloramphenicol and 6 μg ml^−1^ spectinomycin, and placed under low indirect light (30 °C constant light 50 μmol photons m^−2^ s ^−1^ under 1% CO_2_). To clear the plates, additional chloramphenicol and spectinomycin was supplemented underneath the agar to the final concentration of 8 μg ml^−1^ and 16 μg ml^−1^, respectively. Colonies appeared after 10-30 days after the antibiotic addition, which were then streaked on secondary plates for verification. Colony PCR was used to confirm the presence of the expression plasmid as decribed earlier (Venero, et al., 2026) using the primers pDFlac_seq_For 5’-GTTGACTTGTGAGCGGATAACAATGATACTTA-3’ and pDFlac_seq_Rev 5’-CCGCTTCTGCGTTCTGATTTAATCTG-3’. To store the generated strains at -80°C, positive clones were grown in liquid BG-11 to OD_750_=1, pelleted (3min 3000 g) and resuspended gently in 1:7 volume in BG-11 containing 7.5 % DMSO.

### Flv3 immunoblot analysis

For Western blot analysis the *Synechocystis* total protein extracts were prepared and analysed as described earlier (Vuorio, et al., 2021) with slight modifications. The cells from 25 ml culture samples were pelleted and washed with ice-cold washing buffer (50 mM HEPES-NaOH pH 7.5, 30 mM CaCl_2_), centrifuged at 6000g for 8 min at +4°C and subsequently suspended in resuspension buffer (50 mM Hepes-NaOH pH 7.5, 30 mM CaCl_2_, 800 mM sorbitol, 1 mM ε-amino-n-caproic acid). The total protein extracts were obtained by breaking the cells with Bullet Blender Storm 24 at 4°C in the presence of 200 mg of Zirconium oxide 0.15 mm beads. The protein content of the samples was determined using the DC protein assay kit (Bio-Rad). The protein samples were solubilized in Laemmli buffer (Laemmli, 1970) at room temperature and separated electrophoretically on 10% polyacrylamide gels with 6 M urea. The proteins were transferred onto a PVDF membrane (Immobilon-P, Millipore) and immunodetected/detected by a specific antibody for Flv3 (Zhang, et al., 2009) using enhanced chemiluminescence (ECL) (Amersham).

### Sucrose quantitation from culture medium

The samples for sucrose quantitation were harvested from four parallel cultures at 24-hour intervals over a 10-day period, and stored at -80 °C until analysis. The sucrose concentrations in the supernatant were quantified using the Sucrose/d-Glucose Assay Kit (Megazyme, US) based on spectrophotometric analysis of d-glucose released during the enzymatic hydrolysis of sucrose by β-fructosidase, coupled to glucose oxidase/peroxidase GOPOD assay. The reactions were monitored at 510 nm on 96-well plates using microplate reader (Tecan Infinite M200 PRO, CH) with known sucrose standards as reference.

### Proteomic analysis

The cell samples for proteomic analysis were collected from four parallel cultures at three time points (1.5d, 5.5d, 9.5d), harvested by centrifugation and stored at -80 °C until sample preparation. The isolation and digestion of proteins was made according to protocol described earlier (Huokko, et al., 2019) (Vuorijoki, et al., 2016). Briefly, the collected cells were lysed using buffer containing 6M urea in 0.1M Tris-NaOH pH 8 buffer with 1% RapiGest (Waters), 1% PMSF, and an equal volume of glass beads in bead beater. The volume of the crude lysate corresponding to 100 µg of protein was then subjected to reduction with DTT, and alkylation with IAA, followed by precipitation of proteins in cold acetone/ethanol mixture in -20°C overnight. The proteins were then digested with trypsin in 0.05 M Tris -NaOH buffer, pH 8 for 20 h. The mixture of peptides was desalted via Sep-Pack C18 (Waters) columns with the protocol recommended by the manufacturer. Immediately before injection, the samples were spiked with iRT synthetic peptides (Biognosys). Data independent analysia (DIA) for proteomic samples was conducted using a nanoflow HPLC system (Easy-nLC 2000, Thermo Fisher Scientific) interfaced with a Lumos Orbitrap mass spectrometer (Thermo Fisher Scientific) equipped with a high-field asymmetric waveform ion mobility spectrometry (FAIMS Pro) interface in an ionization source, as described in (Lempiäinen, et al., 2025). The acquired raw data were searched with Direct-DIA algorithm in Spectronaut (version 16.1) software against customized FASTA file containing sequences from Cyanobase, iRT and CscB from *E.coli* sequences. The Anova statistics was applied to search for suitable differentiating candidates in pair-wise comparisons. The original raw data from DIA are deposited and publicly available in PRIDE Archive (Vizcaíno, et al., 2016) database with ID PXD080032.

### Metabolomic analysis

The cells for metabolic analysis (OD-normalized) were collected alongside the proteomic samples from four parallel cultures at the given time points, harvested by centrifugation and stored at -80°C until sample preparation. The samples were transported on dry ice to the FIMMS Facility at the University of Helsinki for analysis using a Thermo Vanquish UHPLC+ system equipped with a SeQuant ZIC-pHILIC column (2.1 × 100 mm, 5 µm particle size). This system was coupled to a Q-Exactive mass spectrometer, as detailed by (Muth-Pawlak, et al., 2024).

### Quantitation of intracellular glycogen and PHB

Glycogen and PHB were quantitated from cell aliquots (OD-normalized) collected in parallel to the omics samples. Glycogen analysis was performed using the Total Starch Assay Kit from Megazyme (US), which is based on a spectrophotometric method that quantifies d-glucose generated from the enzymatic hydrolysis of intracellular glycogen coupled with glucose oxidase/peroxidase (GOPOD). Samples were measured in 96-well plates at 510 nm, with glucose acting as the quantitative reference. PHB was analyzed using the D-3-Hydroxybutyric Acid Assay Kit (Megazyme, US), which uses an alkaline lysis method to release D-3-hydroxybutyrate from intracellular PHB. This is followed by a 3-hydroxybutyrate dehydrogenase reaction, which produces a stoichiometric amount of NADH. This was coupled with a diaphorase-catalyzed colorimetric reaction, which was measured on 96-well plates at 492 nm using 3-hydroxybutyrate as the quantitative reference.

### Membrane inlet mass spectrometry (MIMS) gas flux analysis

The steady-state gas flux rates of cell strains were determined by MIMS at increasing light intensities (0, 200 and 500 μmol photons m^−2^s^−1^). These measurements were conducted at atmospheric pressure in a gas-liquid cuvette containing a Teflon membrane (Hansatech Instruments, UK) that separated the cell suspension from the high vacuum line of a Sentinel-PRO magnetic sector mass spectrometer (Thermo Fisher Scientific, USA). The target ions scanned in the mass analysis were m/z 32, 36, and 44, corresponding to ^16^O_2_, ^18^O_2_ and CO_2_, respectively. A remote halogen lamp unit (Dolan-Jenner Industries, USA) calibrated to the required irradiance (Li-250A, LI-COR instruments, USA) was used as the light source. The analytical samples (1 ml) were prepared in a buffer solution corresponding to the culture medium (BG-11, pH 8.0, 50 mM HEPES, 400 mM NaCl). The samples contained the equivalent of 10 μg of chlorophyll. Freshly prepared NaHCO_3_ was added to the reaction medium to provide a CO_2_ reservoir and to prevent carbon limitation during photosynthetic carbon fixation, bringing the final concentration to 1.5 mM. The samples were equilibrated at 25°C in the dark, and purged with N_2_ gas to reduce the amount of ^16^O_2_ signal. Prior to data acquisition, a bubble of ^18^O_2_ (Cambridge Isotope Laboratories Inc., UK) was introduced into the liquid for 2-3 minutes with continuous stirring to equalize the isotope concentrations. Each run was repeated with four culture replicates prepared independently. After primary data processing, the replicates were used to calculate the gas fluxes using the previously described equations (Beckmann, et al., 2009).

## RESULTS

### Engineering the sucrose-producing *Synechocystis* Flv3 overexpression strains

To generate sucrose-excreting *Synechocystis* strains that overexpress the flavodiiron protein Flv3, the existing strain S02 (Thiel, et al., 2019) (Muth-Pawlak, et al., 2024) was engineered further by introducing *flv3* gene as part of the inducible expression plasmid pDF-lac2-cscB-sps-CmR. To ensure high expression levels of Flv3, four different variations of the expression plasmid were constructed, using alternative translational control elements (S3, S4, S7 and A; **Table 1**) upstream the *flv3* coding sequence. The control elements were selected from an established RBS library and have been previously used for dynamic expression of various target proteins in *Synechocystis* (Thiel, et al., 2018). Besides Flv3, the resulting *Synechocystis* strains, named S02oeFlv3 (**Table 1**), overexpressed two additional enzymes, sucrose permease CscB (*Escherichia coli*) and sucrose phosphate synthase SPS (*Synechocystis*) in the glucosylglycerol-phosphate synthase Δ*ggps* deletion background (Thiel, et al., 2019) (**Table 1)**. Throughout the following experiments, S02oeFlv3_S3_, S02oeFlv3_S4_, S02oeFlv3_S7_ and S02oeFlv3_A_ strains were compared against the reference strain S02 that expresses the endogenous Flv3 solely under native regulation.

### Overexpression of Flv3 enhances growth of the sucrose-producing strains under high CO_2_ and high light

In order to compare the growth characteristics of the generated Flv3 overexpression strains, the cells were cultured under 1% CO_2_, 200 μmol photons m^-2^ s ^-1^ continuous white light in BG11 in the presence of 400mM NaCl. While the most significant effects of Flv3 deletion were previously observed in the sucrose-producing strain under these specific conditions (Thiel, et al., 2019), the high CO_2_ and high light are also of general interest from the viewpoint of photobiological bioprocess development. The recorded OD profiles showed systematically higher growth for all the four S02oeFlv3 strains as compared to S02 when cultured in 100ml culture volume in 250ml flasks (**Figure 2**). Interestingly, S02oeFlv3_S4_ grew clearly faster than the rest of the strains throughout the batch culture, reaching OD_750nm_ 8 as compared to OD_750nm_ 6 recorded for S02 over the12-day cultivation period (**Figure 2**). When the experiment was repeated under the same conditions but in 35ml medium volume in 100ml flasks, the cell growth was systematically enhanced and the strain-specific differences were no longer as apparent, apart from S02oeFlv3_S4_ which again grew faster than the other strains (**Supplementary Figure S2**).

**Figure 1:**
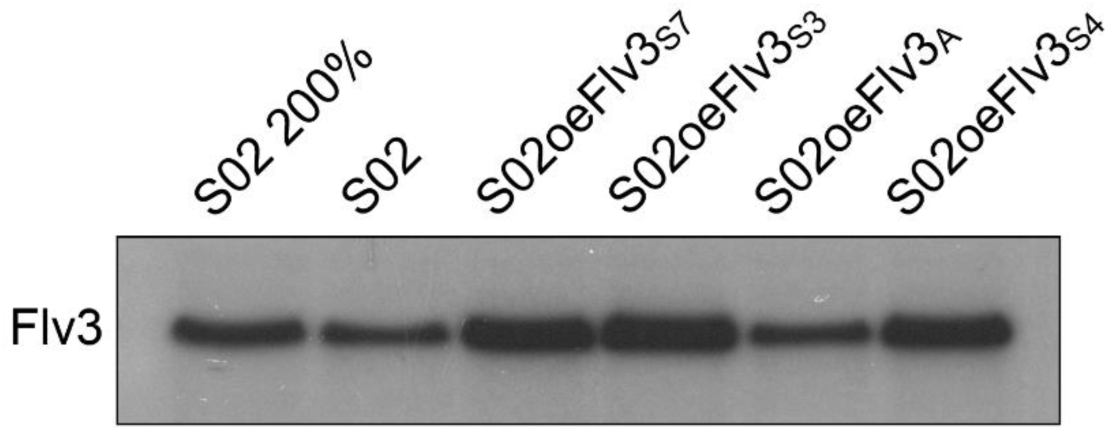
**Immunoblot analysis** of the four alternative *Synechocystis* S02oeFlv3 strains overexpressing Flv3 and the control strain S02 using a Flv3-specific antibody. Protein extracts from culture samples collected at 72h time point were loaded on the gel using 15 µg total protein per well (100%) and separated by SDS-PAGE for the immunoblot analysis. Loading control is shown in **Supplementary Figure S1**.

**Figure 2:**
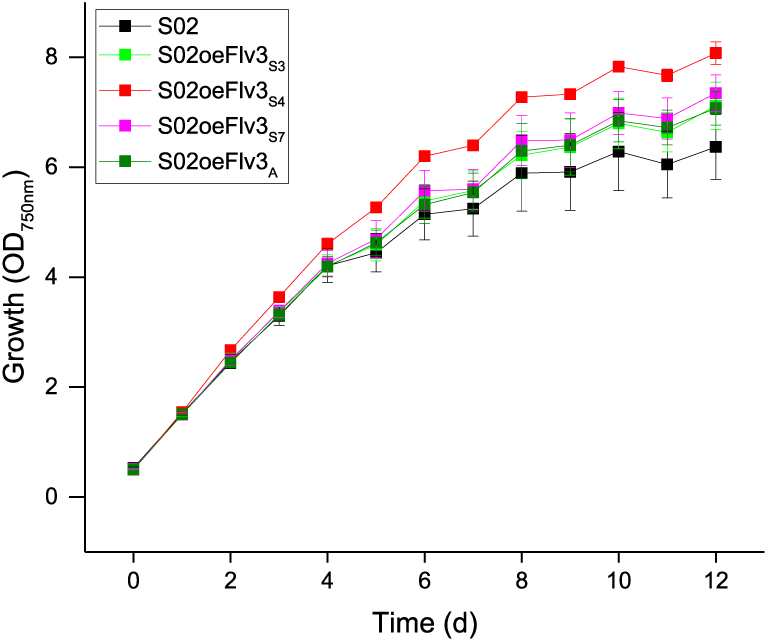
Growth of the four generated Flv3 overexpression strains against S02 in a 12-day batch culture. The cells were grown in 100ml culture volume in 250ml flasks under 1% CO_2_, 200 μmol photons m^-2^ s ^-1^ continuous white light in the presence of 400mM NaCl. The cell growth was monitored spectrophotometrically as optical density at 750nm (OD_750_) at one-day intervals. The averages and standard deviations represent three independent culture replicates (n=3).

### The Flv3 overexpression strains accumulate higher levels of sucrose in the culture medium

The four sucrose-secreting Flv3 overexpression strains were subsequently compared with S02 with respect to sucrose production under the same conditions over a 10-day culture period. The total amount of sucrose that accumulated in the culture medium increased systematically for all the Flv3 overexpression strains (**Figure 3**), corresponding to sucrose concentrations that were ∼100% higher for the most prominent strain S02oeFlv3_S4_ (∼1200mgL^-1^ at d10) as compared to S02 (600mgL^-1^ at d10). While the increase in total sucrose levels was directly linked to faster growth and a higher cell count in the S02oeFlv3_A_ and S02oeFlv3_S3_ cultures, OD-normalized profiles showed enhanced sucrose production per cell in S02oeFlv3_S7_ and, in S02oeFlv3_S4_ (**Figure 3b).** Interestingly, sucrose production per OD reached a maximum in S02 and remained constant after day 2, whereas in S02oeFlv3_S4_ it steadily increased over the next 6 days until d8 (**Figure 3b)**. Consequently, over the eight-day batch cultivation period, the S02oeFlv3_S4_ cells produced over 50% more sucrose per OD than S02 (**Figure 3b**), corresponding to approximately 150mgL^-1^OD_750_^-1^ and 100mgL^-1^OD_750_^-1^ sucrose, respectively.

**Figure 3:**
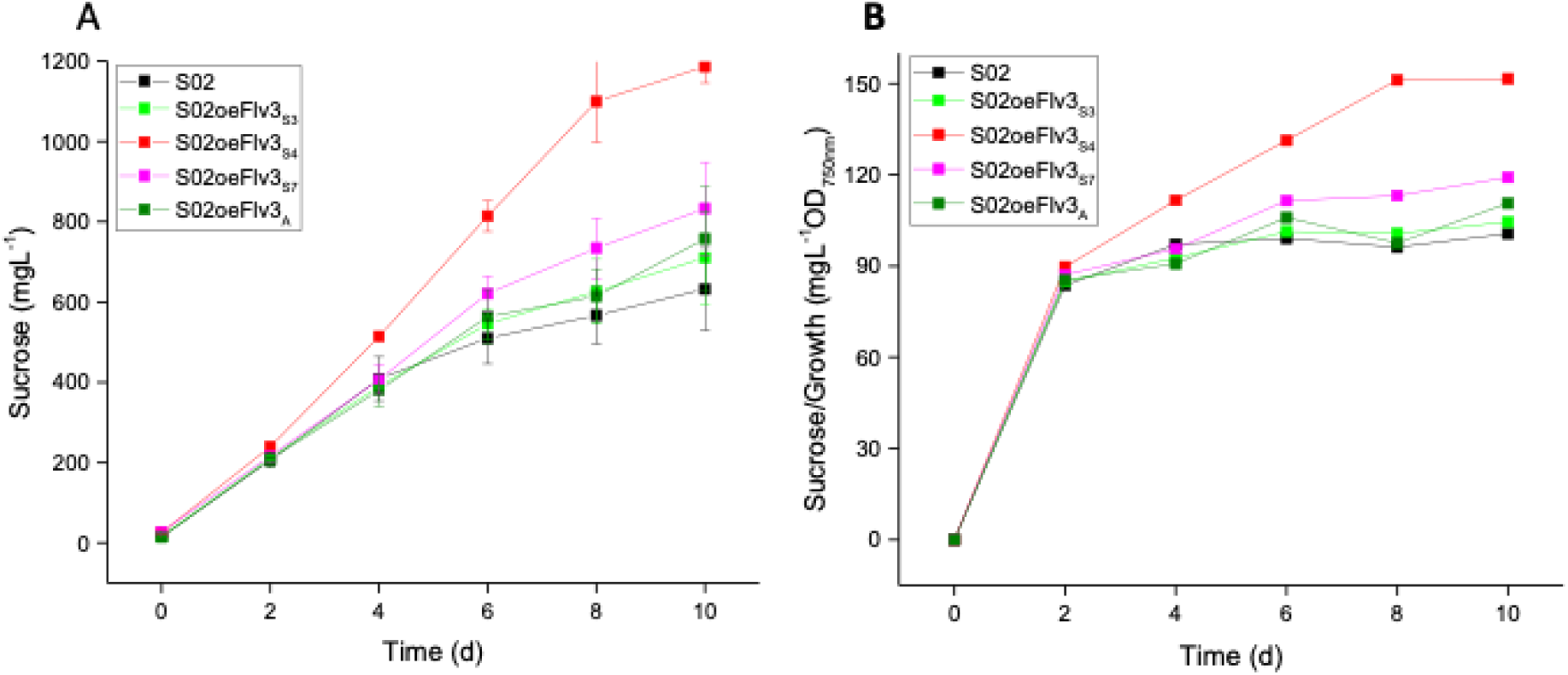
Sucrose production of the four generated Flv3 overexpression strains against S02. **A)** The total concentration of accumulated sucrose and **B)** sucrose concentration normalized to the cell density OD_750_ in a 10-day batch culture. The cells were grown in 100ml culture volume in 250ml flasks under 1% CO_2_, 200 μmol photons m^-2^ s ^-1^ continuous white light in the presence of 400mM NaCl. The averages and standard deviations represent three independent culture replicates (n=3).

### Modulation of growth and sucrose production of S02oeFlv3_S4_

The Flv3 overexpression strain S02oeFlv3_S4_ (referred to as S02oeFlv3 from here on) was selected for the subsequent comparative studies (MIMS, proteomics and metabolite profiling) due to its fastest growth rate (**Figure 2**) and most efficient sucrose production (**Figure 3**). To specify the sampling points and collect the samples, the 12-day main cultivation was repeated for S02 and S02oeFlv3 in four independent replicates. The cultures were grown under 1% CO_2_, 200 μmol photons m^-2^ s ^-1^ continuous white light, and 400mM NaCl in 100ml culture volume using 250ml flasks. The process was monitored by measuring cell growth and sucrose accumulation at 24h intervals for one replicate of each strain (**Figure 4)**. Based on these profiles, the samples were collected from all four parallel S02 and S02oeFlv3 cultures at three different time points, frozen in liquid nitrogen, and stored at -80°C. At the first sampling point (d1.5), the two strains were in the same initial growth phase with no clear phenotypic differences. At the second sampling point (d5.5), the strains were in the middle of the exponential growth phase, with S02oeFlv3 growing faster and producing sucrose at a higher rate than S02. At the third sampling point (d9.5), growth and sucrose accumulation reached a plateau, and the difference in cell density and sucrose concentration between the strains was at its maximum.

**Figure 4:**
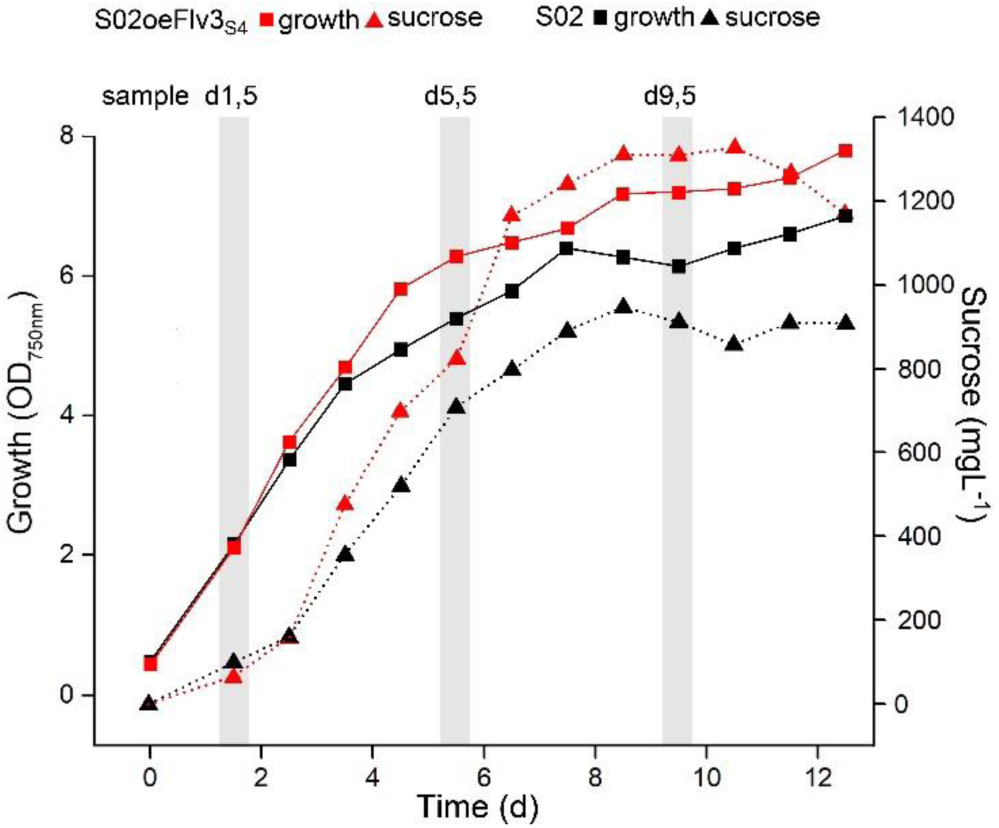
Growth and sucrose production profiles of S02oeFlv3 (red) and S02 (black) showing the sampling points for subsequent analysis. The cells were grown in 100ml culture volume in 250ml flasks under 1% CO_2_, 200 μmol photons m^-2^ s ^-1^ continuous white light in the presence of 400mM NaCl. The cell growth (black squares, solid lines) was monitored spectrophotometrically as optical density at 750nm (OD_750_) at one-day intervals. Sucrose accumulation (red triangles, dotted line) was measured from the culture medium at two-day intervals using a commercial quantitative kit. The vertical shaded bars indicate the sampling points at d1.5, d5.5 and d9.5. The profiles represent single representatives of four independent parallel cultures for each strain.

### Flv3 overexpression strain S02oeFlv3 has increased levels of glycogen and polyhydroxybutyrate

The samples collected for the omics analyses were also used to monitor intracellular changes in accumulation of storage compounds between S02oeFlv3 and S02 during batch cultivation. The glycogen content remained relatively steady in S02 throughout the cultivation process, whereas it increased nearly five-fold in the Flv3 overexpression strain S02oeFlv3 over the eight days between d1.5–d9.5 (**Figure 5a**). Similarly, the accumulation of polyhydroxybutyrate (PHB) was clearly higher in S02oeFlv3 than in S02, although it was observed in both strains, with a final difference of approximately 30% at d9.5 (**Figure 5b**).

**Figure 5:**
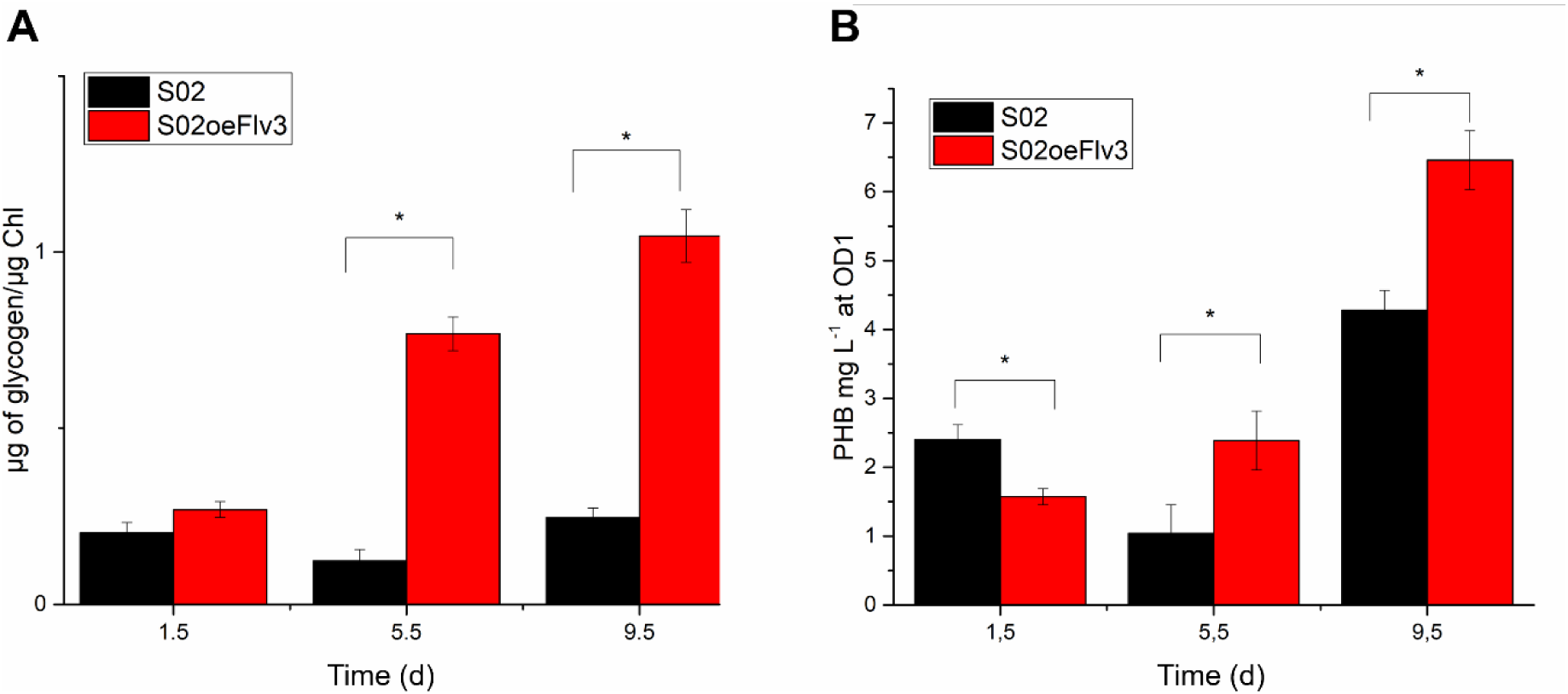
The accumulation of intracellular storage compounds A) glycogen and B) PHB in S02oeFlv3 (black) and S02 (red). The cells were grown in 100ml culture volume in 250ml flasks under 1% CO_2_, 200 μmol photons m^-2^ s ^-1^ continuous white light in the presence of 400mM NaCl, and the samples for storage compound quantitation collected at three time points, 1.5d, 5.5d and 9.5d. The averages and standard deviations represent four independent culture replicates (n=4)

### Gas flux analysis by MIMS revealed increased O_2_ exchange and CO_2_ fixation in Flv3 overexpression strain

Differences in the CO_2_ and O_2_ gas fluxes between the Flv3 overexpression strain S02oeFlv3 and the control strain S02 were addressed by measurements with MIMS (see Materials and Methods; **Figure 6**). The strains were grown in three replicates under 1 % CO_2_ and samples were collected at d1.5, d5.5 and d9.5 (**Figure 4**), and the gas exchange was followed first for 1.5 min in darkness, then 2 min at 200 μmol photons m^-2^ s^-1^ followed by 2 min at 500 μmol photons m^-2^ s^-1^ before turning the lights off. The analysis showed that at the initial sampling point d1.5, the PSII oxygen evolution rates (^16^O_2_ evolution), the oxygen uptake rates corresponding to O_2_ photoreduction and respiration (^18^O_2_ consumption), and the CO_2_ fixation were very similar between the S02oeFlv3 and control S02 strains (**Figure 6 A-B**). Since the O_2_ uptake in the dark (respiratory ^18^O_2_ consumption) was minimal in our experiments, the ^18^O_2_ consumption rates recorded in light can be regarded to mainly represent the O_2_ photoreduction reactions. At the second sampling point d5.5 that represents the exponential growth phase where the phenotypic difference between the strains was most pronounced, the PSII oxygen evolution rates (**Figure 6 C-D**; black line), ^18^O_2_ consumption rates (**Figure 6 C-D**; red line) and CO_2_ fixation (**Figure 6 C-D**; green line) were all clearly higher in S02oeFlv3 than in S02. At the last sampling point d9.5, after reaching the plateau in growth and productivity, the gas fluxes in general had decreased and there were no apparent differences between the control S02 and the Flv3 overexpression S02oeFlv3 strains (**Figure 6 E-F).**

**Figure 6:**
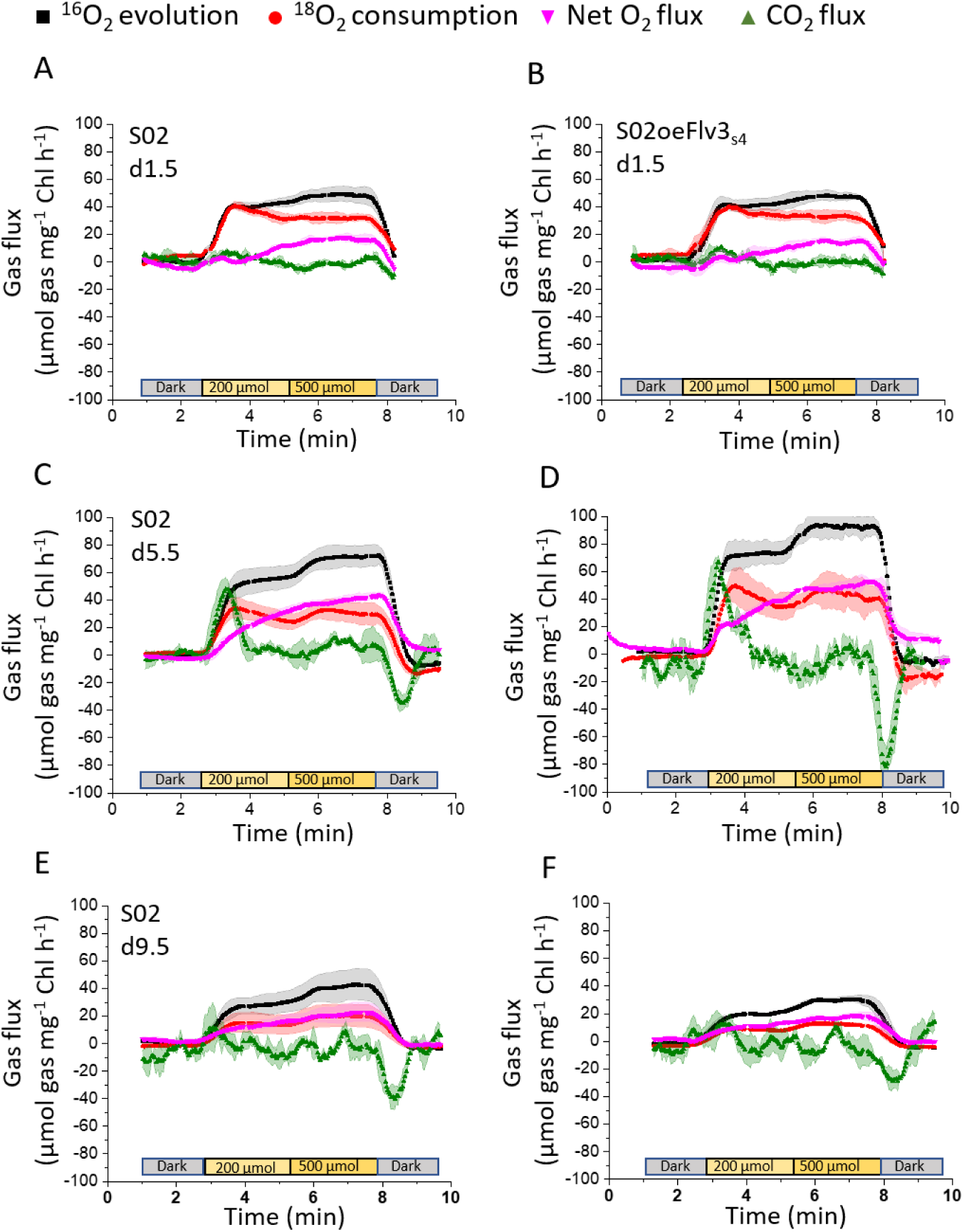
**MIMS gas flux analysis of the sucrose-producing *Synechocystis* Flv3 overexpression strain S02oeFlv3 and the control S02 strain** performed on cells collected at the sampling point **A)** d1.5 occurring at the initial lag phase of growth, **B)** d5.5 at the exponential growth phase and **C)** d9.5 at the final plateau phase. The traces correspond to scanned ion masses m/z 32 for photosynthetic ^16^O_2_ evolution (black), m/z 36 for ^18^O_2_ uptake (red) and m/z 44 for CO_2_ uptake (green). The net oxygen flux is calculated from the O_2_ evolution and uptake rates (purple). The grey bars represent dark incubation before and after each light period, and the yellow correspond to the analysis under 200 μmol photons m^-2^ s^-1^ and 500 μmol photons m^-2^ s^-1^. The measurements were carried out using cells resuspended at concentration equivalent of 10 μg chlorophyll per ml in BG-11 buffered to pH 8.0 with 50 mM HEPES, supplemented with 400 mM NaCl and 1.5 mM NaHCO_3_. Instrument calibrations and gas consumption rates were corrected as per methods outlined (Beckmann, et al., 2009). All curves are an average of minimum three independently analyzed culture replicates (n=3 ±SD).

### The ratio between terminal O_2_ photoreduction and photosynthetic O_2_ evolution is not affected by Flv3 overexpression

Next, we assessed whether the increase in O_2_ photoreduction at d5.5 could be related to Flv3 overproduction in S02oeFlv3, or whether the O_2_ photoreduction rate, catalyzed by canonical Flv1/Flv3 heterodimer, just follows the rate of PSII O_2_ evolution independently of Flv3 overexpression. Notably, the PSII O_2_ evolution rate was at much higher level at d5.5, than at d1.5 or d9.5 in both the S02oeFlv3 and S02 control strains. Calculations of the ratio of the rates of ^18^O_2_ consumption (O_2_ photoreduction) to ^16^O_2_ evolution (the PSII oxygen evolution) from three replicates at three different sampling points and at two different light intensities did not reveal any striking differences in comparisons between the S02oeFlv3 (**Supplementary Figure S3**; orange bars) and S02 control strains (**Supplementary Figure S3**; blue bars). The only noticeable deviation was observed in the higher ratio of the ^18^O_2_ consumption to ^16^O_2_ evolution in the d1.5 samples (around 0.7) compared to the sampling points d5.5 and d9.5 (around 0.4), and this occurred in both S02oeFlv3 and S02. These results clearly demonstrate that the Flv3 protein, which is overexpressed in S02oeFlv3, is not involved in additional O_2_ photoreduction, either as a component of the Flv1/3 heterodimer or as an independent Flv3 homooligomer.

### Proteome analysis revealed clear differences between S02oeFlv3 and S02 strains during sucrose-producing growth

The whole-cell samples from S02 and S02oeFlv3 strains, collected as four independent culture replicates at the three selected time points (d1.5, d5.5 and d9.5), were subjected to total protein isolation and digestion, followed by LC MS/MS DIA analysis. 2863 proteins were identified in direct-DIA approach (**Supplementary Table S1**), which covers 88% of the entire predicted *Synechocystis* proteome (Kaneko and Tabata, 1997). The 24 samples grouped into six clearly independent clusters that represented horizontal separation to S02 and S02oeFlv3 strains and vertical separation to time points of sample collection corroborating high reproducibility of the data (**Figure 7A**). The dispersion of the datapoints on volcano plots (**Figure 7B-D**) reflect the increasing differentiation between two strains in the course of time. The differences are initially, on d1.5, small (52 differentially regulated proteins), with longer cultivation time, more differentiating proteins (222 differentially regulated proteins) were evidenced on d5.5 (Figure 7C) reaching highest differentiation (395 differentially regulated proteins) on day 9.5 (**Figure 7D**).

**Figure 7:**
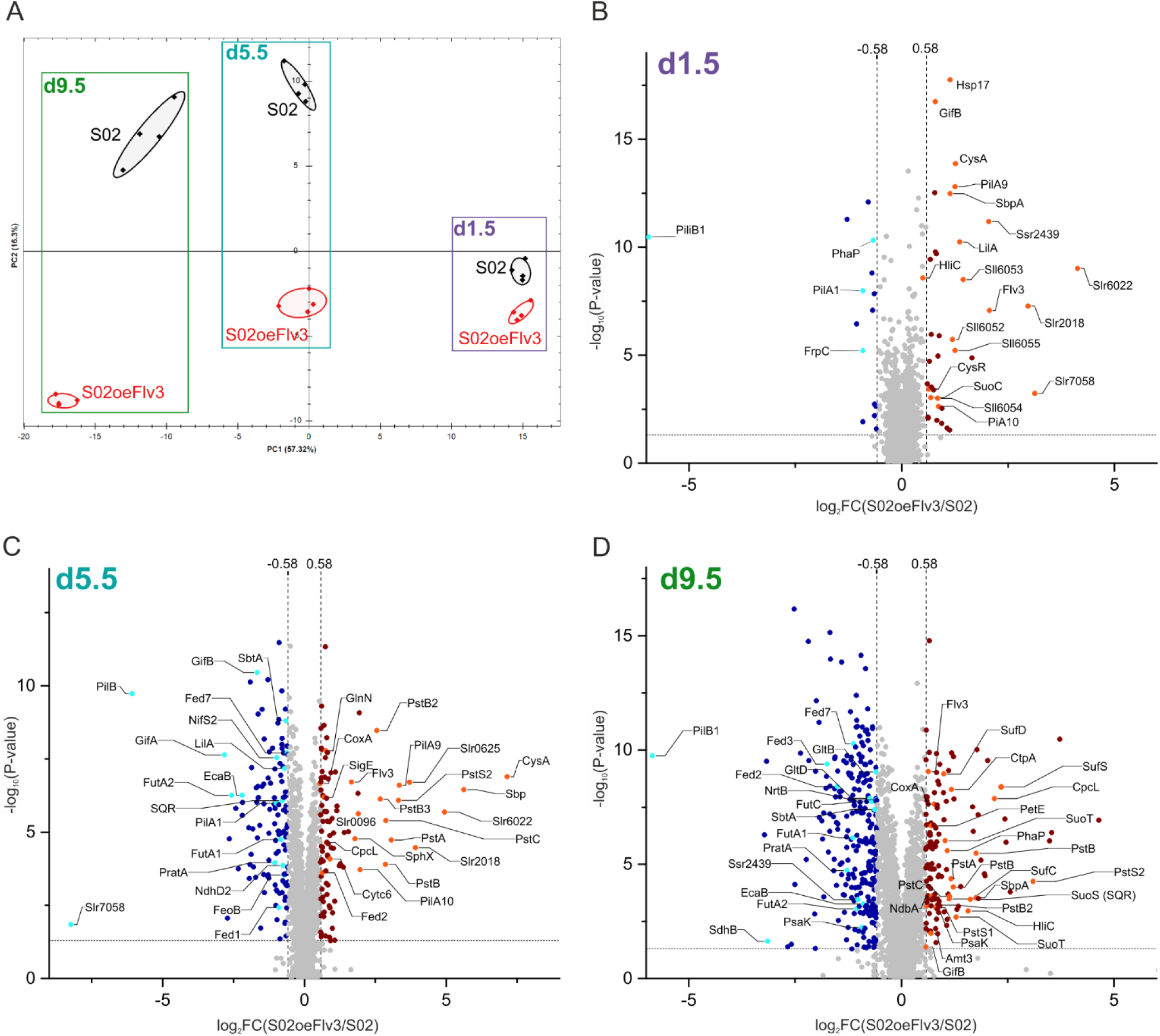
Proteomic comparison between the S02oeFlv3 and S02 strains during the sucrose-producing growth. **A)** Principal Component Analysis (PCA) of the LC-MS/MS data. Volcano plots of the S02oeFlv3 and S02 proteomes at the sampling points **B)** d1.5, **C)** d5.5 and **D)** d9.5 as shown in Figure 4. The blue dots represent significantly downregulated proteins and brown dots significantly upregulated proteins in S02oeFlv3 as compared to S02. Light blue and orange dots represent labeled down and up regulated proteins, respectively. See **Table 2** and **Supplementary Table S1** for details of individual protein expression in the S02oeFlv3 and S02 strains.

**Table 2:** Quantitative protein-level differences observed in the S02oeFlv3 strain in relation to S02 at the three sampling points d1.5, d5.5 and d9.5 during the sucrose-producing growth. The comparison is presented as a log_2_ fold change (FC) with red font indicating upregulation and blue font indicating downregulation in S02oeFlv3 compared to S02. Statistically significant changes (*p* < 0.05 or adjusted) are shown in **bold**.

| ORF | Prot ID | Log<sub>2</sub> FC S02oeFlv3/S02 |  |  | Protein function |
| --- | --- | --- | --- | --- | --- |
|  |  | d1.5 | d5.5 | d9.5 |  |
| **sII0550** | Flv3 | **2.1** | **1.7** | **0.6** | Photosynthesis and respiration |
| **sII1521** | Flv1 | **0.2** | **0.3** | **0.4** | Photosynthesis and respiration |
| **ssr2439** | Ssr2439 | **2.0** | - | **-1.0** | Sulfate transporter operon |
| **slr1455** | CysA | **1.3** | **7.1** | **1.3** | Sulfate transporter operon |
| **slr1452** | SbpA | **1.1** | **5.6** | **1.8** | Sulfate transporter operon |
| **slr0096** | Slr0096 | **0.3** | **1.9** | **-0.1** | Sulfate permease (hypothetical) |
| **slr1544** | LilA | **1.4** | **-0.7** | -0.6 | Small CAB-like Protein (SCP) |
| **ssl1633** | HliC | **0.5** | **0.2** | **1.6** | Small CAB-like Protein (SCP) |
| **sII0680** | PstS1 | **0.4** | **1.0** | **0.8** | Phosphate uptake |
| **sII0681** | PstC | **-0.1** | **2.9** | **1.1** | Phosphate uptake |
| **sII0682** | PstA | **0.2** | **3.1** | **1.2** | Phosphate uptake |
| **sII0683** | PstB | **0.2** | **2.9** | **1.2** | Phosphate uptake |
| **sII0684** | PstB2 | -0.1 | **2.6** | **0.8** | Phosphate uptake |
| **slr1247** | PstS2 | **0.4** | **3.3** | **3.1** | Phosphate uptake |
| **slr1250** | PstB3 | 0.1 | **2.7** | **1.8** | Phosphate uptake |
| **ssl0020** | Fed1 | **-0.3** | **-0.9** | 0.3 | Photosynthesis and respiration |
| **sII1382** | Fed2 | 0.1 | **0.6** | **-1.5** | Photosynthesis and respiration |
| **slr1828** | Fed3 | 0.0 | **0.3** | **-1.7** | Photosynthesis and respiration |
| **sII0662** | Fed7 | 0.0 | **-0.6** | **-1.1** | Photosynthesis and respiration |
| **slr1137** | CoxA | 0.0 | **0.6** | **0.8** | Photosynthesis and respiration |
| **slr0851** | NdbA | -0.1 | **0.1** | **0.7** | NDH-2 |
| **slr1291** | NdhD2 | **-0.5** | **-0.8** | **0.4** | Cyclic electron transfer |
| **sII1796** | PetJ | **0.1** | **0.9** | **0.5** | Soluble electron carrier |
| **slI0199** | PetE | 0.1 | **-0.2** | **0.7** | Soluble electron carrier |
| **slr2048** | PratA | **0.3** | **-1.0** | **-1.3** | PSII assembly factor |
| **slr0008** | CtpA | 0.0 | **0.2** | **1.2** | PSII assembly factor - D1 processing |
| **slr1295** | FutA1 | -0.1 | **-0.8** | **-1.1** | Iron transport |
| **slr0513** | FutA2 | **0.3** | **-2.6** | **-1.0** | Iron transport |
| **slI1878** | FutC | -0.1 | **-0.5** | **-0.7** | Iron transport |
| **ssr0390** | PsaK1 | **-0.2** | -0.1 | **0.6** | PSI |
| **slI0629** | PsaK2 | -0.2 | **-0.3** | **-0.9** | PSI |
| **slI1471** | CpcL | 0.1 | **0.8** | **2.2** | PSI-associated PBS linker |
| **slI1330** | Rre37 | 0.0 | **0.4** | **0.6** | RRE CCM regulator |
| **slr0051** | EcaB | **0.2** | **-2.2** | **-0.9** | CCM |
| **slr1512** | SbtA | 0.0 | **-0.7** | **-0.6** | CCM |
| **slI1027** | GltD | **0.0** | **0.3** | **-0.7** | Glutamate synthase |
| **slI1502** | GltB | 0.0 | **0.2** | **-0.6** | Glutamate synthase |
| **slr0288** | GlnN | **-0.2** | **0.7** | **0.4** | Glutamine synthetase |
| **ssl1911** | GifA | - | **-2.8** | 0.0 | GS-GoGat regulator |
| **slI1515** | GifB | **0.8** | **-1.7** | **0.6** | GS-GoGat-regulator |
| **slI1689** | SigE | **0.4** | **0.7** | **-0.5** | Sigma factor |
| **slI1455** | NarM | 0.1 | 0.1 | **-0.6** | Nitrate assimilation |
| **slI1450** | NrtA | **-0.2** | 0.1 | **-0.2** | Nitrate assimilation |
| **slI1451** | NrtB | **-0.4** | **0.2** | **-0.6** | Nitrate assimilation |
| **slI1452** | NrtC | -0.1 | **0.1** | **-0.4** | Nitrate assimilation |
| **slI1453** | NrtD | -0.1 | **0.2** | **-0.3** | Nitrate assimilation |
| **slI0537** | Amt3 | 0.0 | **0.5** | **0.7** | Ammonium assimilation |
| **slI5035** | SuoR | **0.3** | **-0.3** | **1.0** | SuoRSCT operon |
| **slI5036** | SuoS | **0.0** | **-0.8** | **1.6** | SuoRSCT operon/SQR |
| **slr5037** | SuoC | **0.7** | **-0.6** | **0.4** | SuoRSCT operon |
| **slr5038** | SuoT | **0.0** | **-0.2** | **1.3** | SuoRSCT operon |
| **ssl2501** | PhaP | **-0.7** | **0.9** | **1.1** | PHB metabolism |
| **P30000** | CscB | **-0.2** | **0.3** | **0.1** | Sucrose transporter (*E.coli*) |
| **slr0076** | SufD | 0.0 | **0.2** | **1.0** | Fe-S cluster biosynthesis |
| **slr0075** | SufC | 0.1 | **0.1** | **1.1** | Fe-S cluster biosynthesis |
| **slr0077** | SufS | 0.0 | **0.3** | **2.3** | Fe-S cluster biosynthesis |
| **slr0625** | Slr0625 | 0.0 | **3.7** | **3.7** | sodium dependent glutamate transporter |

### Only few protein-level differences between S02oeFlv3 and S02 were evident in the beginning of the sucrose-producing growth, at d1.5

The overexpression of Flv3 in S02 background resulted in 4-fold increase in the abundance of the Flv3 protein (log_2_FC=2.07) in S02oeFlv3, which subsequently reduced to 300% excess at d5.5 and 50% excess at d9.5. The increase in Flv3 at d1.5 was not accompanied by any phenotypic changes in growth or sucrose production, and only a few other proteins showed significant expression differences between S02 and S02oeFlv3. Differentially expressed proteins included upregulated small chlorophyll (SCP) binding proteins LilA and HliC in S02oeFlv3. Also, proteins from the sbpA-cysTWA operon (Lee, et al., 2021) involved in sulfate binding (SbpA) and transport (CysA) were upregulated in S02oeFlv3, and remained at high level throughout the growth period. Another component of the operon Ssr2439, hypothetically performing a regulatory role, was highly induced on d1.5 but declined by d9.5. Notably, the abundance of the Flv1 protein in S02oeFlv3 remained at similar level with that in the S02 control strain throughout the sucrose-producing growth period (**Table 2**). Similarly, the sucrose pathway proteins overexpressed in both S02 and S02oeFlv3 (**Table 1**) did not reveal any differential expression between the strains (**Table 2**).

### Differences in proteome expression between S02 and S02oeFlv3 became more prominent at sampling points d5.5 and d9.5

Thylakoid membrane-related proteins, such as PSII assembly protein PratA and LilA decreased in abundance towards the end of the culture period in S02oeFlv3 compared to S02. On the contrary, the levels of HliC (He, et al., 2001) involved in high-light photoprotection, and the D1 processing protein CtpA increased in S02oeFlv3 as observed at d9.5. Transient downregulation of ferredoxin 1 (Fed1) was detected in S02oeFlv3 strain at d5.5 while Fed2, Fed3 and Fed7 decreased in abundance only later at d9.5. The soluble electron carriers transferring electrons from cytochrome *b_6_f* (Cyt *b_6_f*) to PSI, such as cytochrome c6 (PetJ) showed upregulation at d5.5, while plastocyanin (PetE) increased in abundance in S02oeFlv3 by d9.5. PSI-related proteins, on the other hand, involved an increase in the phycobilisome linker protein (CpcL) at d5.5 and d9.5 as well as in the PSI protein PsaK1, while the PsaK2 isoform decreased in abundance at d9.5 in S02oeFlv3. In addition, a specific NdhD2 subunit of the NDH-1_2_ complex, which is responsible for cyclic (CET) and respiratory electron transfer, increased in abundance at d9.5 in the S02oeFlv3 strain. Other signs of increased respiratory activity at d9.5 in S02oeFlv3 compared to S02, included an increased abundance of NdbA that receives electrons from metabolically reduced NADH and transfers them to the PQ pool, together with the accumulation of the cytochrome c oxidase (CoxA) subunit of the terminal respiratory complex cytochrome c oxidase (Cox).

Apart from thylakoid-related proteins, also other redox proteins showed differential expression between S02oeFlv3 and S02 during the sucrose production experiment. Intriguingly, three out of four proteins encoded by the SuoRSCT operon, located in the native plasmid pSYSM, showed upregulation at d9.5. SuoS belongs to sulfide:quinone reductases (SQRs) which is a member of the disulfide oxidoreductase (diSR) flavoprotein family. SuoS has been shown to catalyze light-dependent electron transfer from sulfide group (**-**SH) directly to membrane-bound plastoquinone (PQ) pool as the electron acceptor (Nagy et al., 2014). The other upregulated components of Suo operon in S02oeFlv3 were SuoR that belongs to an ArsR transcriptional regulatory proteins involved in arsenate and sulfide detoxification, SuoC with unknown function and SuoT that was initially assigned as a chromate transporter, but later it has been shown to function as arsenite (AsO_3_^3−^ ) uptake transporter (Nagy, et al., 2014).

Only minor differences were observed between S02oeFlv3 and S02 in protein abundances of central carbon and nitrogen metabolism. Two proteins involved in carbon concentration mechanisms (CCM), the sodium-dependent bicarbonate transporter SbtA and the thylakoid-located carbonic anhydrase EcaB showed decreased abundance from d5.5 onwards in S02oeFlv3. On the other hand, the regulatory proteins SigE (Osanai, et al., 2005) (Osanai, et al., 2013) and Rre37 (Joseph, et al., 2014), which control sugar catabolic genes and stimulate the biosynthesis of polyhydroxybutarate (PHB) and glycogen (Glk), respectively, were significantly upregulated in S02oeFlv3 at d5.5 and d9.5 compared to S02. At the same time, phasin PhaP (Hauf, et al., 2015), which contributes to PHB accumulation by stabilizing the formation of PHB granules in *Synechocystis*, showed increased expression in S02oeFlv3 at d5.5 and d9.5 compared to S02. In parallel, a strong decrease in abundance of GS-GoGat pathway regulators GifA/B and increased abundance of glutamine synthetase GlnN and ammonium transporter Amt3 indicated transient induction of nitrogen assimilation at d5.5. However, at d9.5 the extent of nitrogen assimilation had decreased in S02oeFlv3, as deduced from increased abundance of GifB regulator and lowered relative levels of ferredoxin-dependent glutamate synthase (GltB and GltD), and NarM, regulating nitrate reductase activity. This indicates that S02oeFlv3 cells sense transient imbalance in the C/N ratio and induce N assimilation but do not upregulate nitrate transporters such as the Nrt system typically expressed under nitrogen depletion conditions as shown in (Muth-Pawlak, et al., 2022).

High energy status of the S02oeFlv3 strain was indicated by induction of phosphate uptake operon PstA-C together with phosphate binding proteins (PstS1/2) and phosphate kinase from d5.5. Iron uptake system FutA1/2 and FutC decreased from d5.5 in S02oeFlv3 strain. In addition, three proteins from SufBCDS-operon, responsible for biosynthesis of the iron-sulfur clusters, were induced in S02oeFlv3 strain at d9.5. These proteins included cysteine desulfurase SufS, SufE that enhances SufS activity, SufC, an ATP-dependent iron transporter and SufD, which, together with SufB and C, forms a scaffold complex where the Fe-S clusters are assembled (Gao, 2020). Increased protection against osmotic stress in S02oeFlv3 strain, compared to S02, was shown by increased abundance of Slr0625 sodium dependent glutamate transporter from d5.5.

### Differential changes in metabolite composition between the S02oeFlv3 and S02 strains during sucrose-producing growth

Quantitative metabolomics analysis using LC–MS/MS was carried out to evaluate alterations in metabolic pathway activities in the S02 and S02oeFlv3 strains during sucrose-producing growth. Metabolite profiling (**Table 3**) revealed an increased accumulation of 2-OG, aconitate and citrate involved in the oxidative part of the TCA cycle (Zhang, et al., 2016) in S02oeFlv3 compared to S02 at d1.5. The abundance of aconitate and 2-OG further increased in S02oeFlv3 until d9.5, while the citrate level stayed at d5.5 level. Pyruvate, an important metabolic precursor derived from glycolytic pathway, was only detected in S02oeFlv3 strain at d5.5 and its level increased until d9.5. At d5.5, most of the amino acids (including serine, alanine, arginine, aspartate, threonine, isoleucine and tryptophan) accumulated at higher levels in S02oeFlv3 than in S02, apparently reflecting S02oeFlv3’s faster growth rate. Metabolites from the reductive part of the TCA cycle, such as malate and fumarate, were less abundant in S02oeFlv3 than in S02 at d5.5 and d9.5. At d5.5, the levels of ATP and oxidized glutathione (GSSG) were also lower in S02oeFlv3 than in S02. The extremely low level of GSSG during the fast growth period at d5.5, points to a highly reduced cellular environment in S02oeFlv3 compared to S02. Lactate and acetate accumulated at d9.5, which may indicate low oxygen availability. The levels of GSSG and NADH increased towards d9.5 in S02oeFlv3 compared to S02.

**Table 3:** Quantitative differences observed in intracellular metabolite levels in the S02oeFlv3 strain in relation to S02 at the three sampling points d1.5, d5.5 and d9.5 during sucrose-producing growth. The comparison is presented as a fold change (FC) with blue font indicating downregulation and red font indicating upregulation in S02oeFlv3 compared to S02. * Pyruvate was detected only in S02oeFlv3 at d5.5 and 9.5. 2-OG; 2-oxoglutarate, 2-PGA; 2-phosphoglycerate, 6-P-Glc; glucose-6-phosphate; AcCoA; acetyl-CoA, Ac-OH; acetate; Acon; aconitate, Ala; alanine, Arg; arginine, Asp; aspartate, Cit; citrate, DHAP; dihydroxyacetone phosphate, Fum; fumarate, Glu; glutamine, GSSG; glutathione disulfide, IsoLeu; isoleucine, LacAc; lactate, Lys; Lysine, MaAc; malate, NADH; nicotinamide adenine dinucleotide (reduced), NADP; nicotinamide adenine dinucleotide phosphate (oxidized), Orn; ornithine, P-enolPyr; phosphoenolpyruvate, Ser; serine, Suc; succinate, Tre; trehalose, Trp; tryptophan; UDP; uridine diphosphate.

| Metabolite | FC S02oeFlv3 vs S02 |  |  |
| --- | --- | --- | --- |
|  | d1.5 | d5.5 | d9.5 |
| 2-OG | 2.4 | 4.7 | 14.5 |
| 2-PGA |  |  | -1.6 |
| 6-P-Glc |  |  | 1.5 |
| AcCoA |  | -1.5 |  |
| AcOH |  |  | 1.9 |
| Acon | 2.2 | 4.4 | 8.3 |
| Ala |  | 2.5 | 1.8 |
| Arg |  | 2.7 | 1.7 |
| Asp |  | 2.0 |  |
| Cit | 1.6 | 3.2 | 3.6 |
| DHAP |  | -1.5 |  |
| Fum |  | -2.4 | -1.9 |
| Glu |  |  | 1.6 |
| GSSG |  | -27.1 | 1.6 |
| IsoLeu |  | 1.9 | 1.5 |
| LacAc |  |  | 2.0 |
| Lys |  | 2.9 | 2.0 |
| MaAc |  | -2.1 | -1.8 |
| NADH |  |  | 2.5 |
| NADP |  |  | -1.6 |
| Orn |  | 5.1 | 2.0 |
| PEP |  |  | -1.6 |
| Ser |  | 4.3 | 2.1 |
| SucAc | 1.5 |  | 1.5 |
| Tre |  | 2.0 |  |
| Trp |  | -1.5 | -1.6 |
| UDP |  | -2.0 | -1.9 |
| Pyruvate* | ND | Only in<br>S02oeFlv3 | Only in<br>S02oeFlv3 |

## DISCUSSION

### *Synechocystis* cells benefit from the overexpression of Flv3

Flavodiiron proteins Flv1 and Flv3 function as Flv1/Flv3 heterodimers in *Synechocystis* and play a key role in regulating photosynthetic and respiratory electron transfer, and consequently the redox homeostasis in cyanobacterial cells. The Flv1/Flv3 heterodimer has been a subject of extensive research, with an assigned role in a Mehler-like reaction, where it directs electrons derived from PSII water-splitting reactions to molecular oxygen (oxygen photoreduction) via LET (Allahverdiyeva, et al., 2013).

In addition to the photoprotective oxygen reduction activity of the Flv1/Flv3 heterodimer, we present new data on the Flv3 protein’s independent function. We used an engineered *Synechocystis* strain that produces and excretes sucrose, and therefore harbours strong electron sinks (Thiel, et al., 2019) as the background to overexpress the Flv3 protein at different levels (the S02oeFlv3 strains). All the four generated S02oeFlv3 strains could be distinguished from the S02 control (with WT-level of Flv3; **Figure 1**) on the basis of enhanced cell growth (**Figure 2**) and sucrose production efficiency (**Figure 3**). The S02oeFlv3_S4_ strain, which expresses Flv3 under the RBS derived from the *psbA2* gene in *Synechocystis* (S02oeFlv3 from here on), exhibited the highest sucrose production and fastest growth in all experiments, and was therefore used for a more in-depth characterization of Flv3 overexpressing cells. The S02oeFlv3 strain also demonstrated enhanced sulphur metabolism. This raises fundamental questions about the potential role of the Flv3 protein in modulating cellular metabolism, which will be discussed later.

### The growth mode of sucrose-producing Synechocystis strains is modulated by drastic differences in Flv3 expression

The first indications of an independent functional role of Flv3 in *Synechocystis*, aside from its role in the Flv1/Flv3 heterodimer, can be inferred by comparing the sucrose-producing control strain S02 with respective *flv3* overexpression (S02oeFlv3) and *flv3* deletion (S02:Δ*flv3)* (Muth-Pawlak, et al., 2024) mutants. Both S02:Δ*flv3* and S02oeFlv3 grow, after an initial short lag phase, faster than the S02 strain. Interestingly, the *flv3* deletion and overexpression mutants, however, adopt completely different growth modes. The S02:Δ*flv3* strain begins to consume the sucrose produced via the engineered sucrose pathway, thereby improving the growth by transition to mixotrophic mode (Muth-Pawlak, et al., 2024). Conversely, the overexpression S02oeFlv3 strain enhances the accumulation of sucrose in the medium compared to S02 (**Figure 3**) and simultaneously improves autotrophic growth (**Figure 2**), as would be expected from the strengthening of the capacity of the electron sink (Abramson, et al., 2016). An increase in growth was also observed in WT *Synechocystis* overexpressing Flv3 when cultured under low-light and elevated CO_2_ (Hasunuma, et al., 2014). Taking into account the putative novel function of the excessive Flv3, it is important to consider the ratio of Flv3 to Flv1 proteins in the cell. In WT *Synechocystis*, Flv3 and Flv1 are encoded by separate genes under independent transcriptional control, and observed in WT to be present in the molar ratio 4:1 (Jackson et al 2023). This suggests that in addition to the Flv1/Flv3 heterodimer (ratio 1:1), an extra pool of Flv3 is always available to form Flv3 homooligomers in the WT, as also expected for the control strain S02 that harbors the intact native genes *flv3* and *flv1*.

When comparing S02, S02:Δ*flv3* and S02oeFlv3 it is important to note that, unlike the other strains, S02:Δ*flv3* lacks not only the putative Flv3 homooligomer but also the canonical Flv1/Flv3 heterodimer. This can be deduced from the fact that neither Flv1 nor Flv3 were detected by mass spectrometry in S02:Δ*flv3* (Muth-Pawlak, et al., 2024). Importantly, such a loss of both Flv3 forms decreases the electron sink capacity and triggers mixotrophic phenotype of S02:Δ*flv3*. In contrast, the S02 control strain and the S02oeFlv3 strain differ in that the latter exhibits much higher Flv3 protein expression (**Figure 1**), while Flv1 protein expression remains the same (**Table 2**). Thus, the superior phenotype of S02oeFlv3 over S02 is solely due to higher levels of the Flv3 protein (**Figure 1**) and its apparent function as a Flv3 homooligomer providing beneficial growth capacity. The clear phenotypic differences observed between the four parallel Flv3 overexpression strains further underline that expression optimization in the target host is essential to take full advantage of Flv3 in strain engineering **Figure 3**, **Table 1**).

### Gas exchange measurements support an independent physiological role for overexpressed Flv3 in *Synechocystis* engineered for enhanced sucrose production

Considering the physiological role of Flv3 homooligomers in cellular context, the gas exchange experiments performed by MIMS (**Figure 6**, **Supplementary Figure S3**) indicate that they are not involved in O_2_ photoreduction. Although the S02oeFlv3 strain showed apparently higher oxygen photoreduction at d5.5 (**Figure 6**), when the sinks were functioning at maximal efficiency (see **Figure 6**), the ratios of ^18^O_2_ uptake to PSII ^16^O_2_ evolution remained similar to S02 (**Supplementary Figure S3**). This indicates that, in defined strains and growth conditions, the increase in PSII oxygen evolution rate is linked to a similar relative increase in O_2_ photoreduction rate (**Supplementary Figure S3**). This phenomenon is also seen in WT *Synechocystis* grown at saturating CO_2_ of 3% (Santana Sanchez, et al., 2019). In this case, increasing the PSII O_2_ evolution rate by elevating the light intensity from 500 to 1000 µmol photons m^-2^ s^-1^ also increased the O_2_ photoreduction rate (recalculated from data form (Santana Sanchez, et al., 2019)), while keeping the ratios of ^18^O_2_ uptake to PSII ^16^O_2_ evolution relatively constant, 0.28+/-0.05 and 0.36+/-0.05, respectively. Taken together, these findings provide evidence for one or more as yet unknown function(s) for putative Flv3/Flv3 homodimers, or other homooligomer states, as was also proposed earlier (Mustila, et al., 2016) (Thiel, et al., 2019) (Eckardt, et al., 2024). In search of Flv3-dependent metabolic pathways that are currently unknown, we make use of engineered *Synechocystis* strains with enhanced sucrose production. These strains act as a saturated photosynthetic electron sink making the PSII oxygen evolution and the LET rates mainly dependent on light intensity (**Figure 8**).

**Figure 8:**
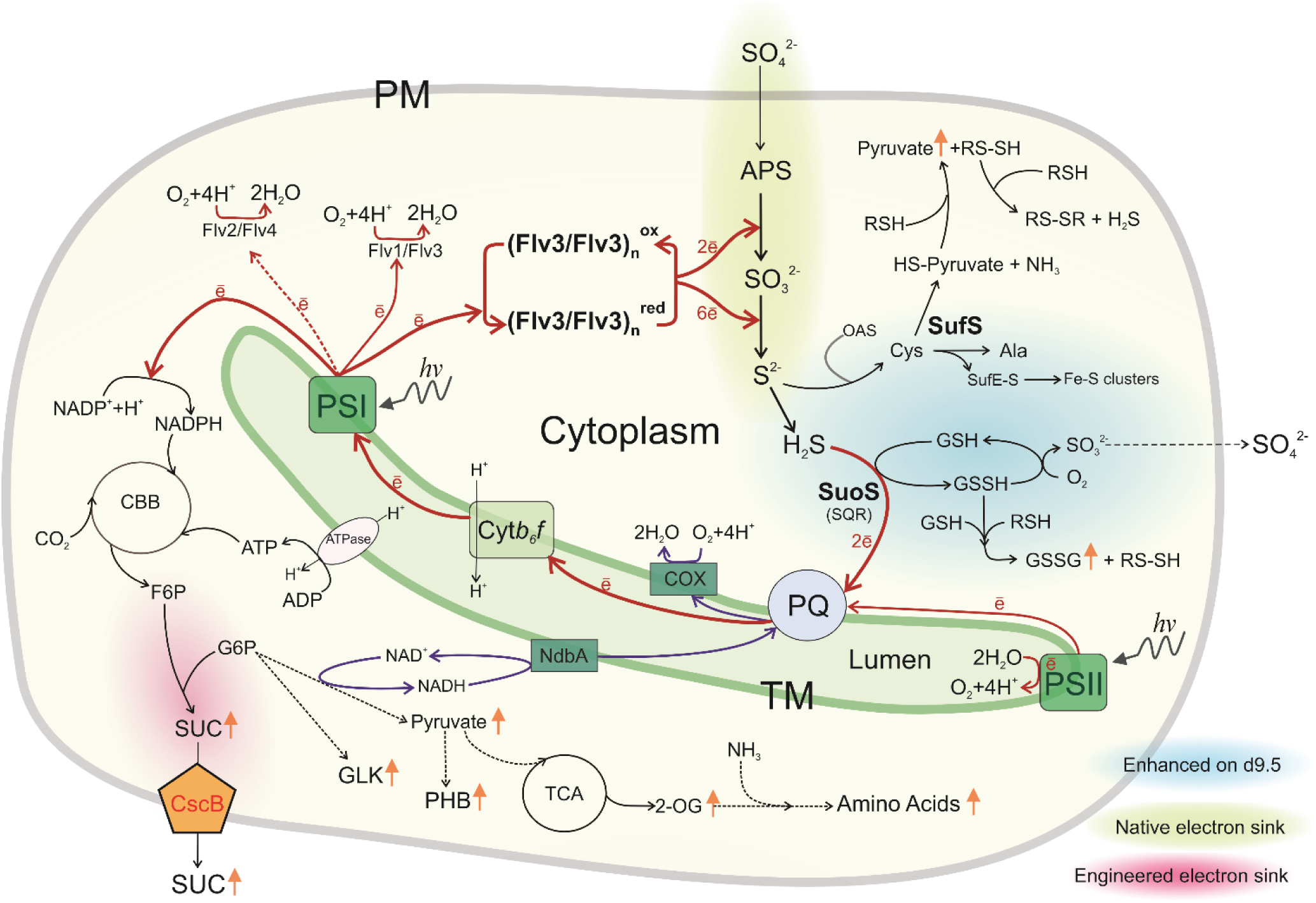
Hypothetical scheme for the function of the Flv3 homooligomer in cellular redox homeostasis and sulfate metabolism in *Synechocystis* sucrose producing strain S02oeFLV3. For details, see the main text. Red arrows indicate photosynthetic electron flow, blue arrows respiratory electron flow and black arrows cellular metabolism. Orange arrow indicates an increase in metabolite abundance during sucrose production. PM - plasma membrane, TM - thylakoid membrane, Flv - flavodiiron protein, APS - adenosine 5’ phosphosulfate, SufS – cyst(e)in desulfuraze, OAS – O-acetyl-serine, Cys – cysteine, GSH – reduced glutathione, GSSG – oxidized glutathione, GSSH – persulfidated glutathione, RSSH – persulfidated molecule PQ - Plastoquinone pool, PSI - photosystem I, PSII - photosystem II, Cytb_6_f - cytochrome b_6_f, SQR - sulfide:quinone reductase, HS-Pyruvate – 3-mercaptopyruvate, CBB -Calvin-Benson-Bassham cycle, SUC- sucrose, GLK – glycogen, PHB – polyhydroxybutarate, ATPase – ATP synthase, CscB – sucrose permease (*E.coli*), F6P – fructose-6-phosphate, G6P – glucose -6-phosphate, 2-OG- 2-oxoglutarate, COX – cytochrome c oxidase, TCA – tricarboxylic acid cycle.

### Proteome and metabolite screening reveal specific features underlying the superior phenotypes of the Flv3 overexpressing sucrose producing *Synechocystis* strain

During the early growth phase (d1.5 lag phase) only minor proteomic differences between the S02 and S02oeFlv3 strain were observed. The most notable distinction was the higher abundance of sulfate transporters in the S02oeFlv3 strain (to be discussed in more detail later). Otherwise, the Flv3 overexpression in S02oeFlv3 revealed only minor phenotypic or molecular-level differences compared to SO2. These included an increase in 2-OG, which is indicative of enhanced glycolytic activity, and the upregulation of proteins that function in photoprotection and the maintenance of the photosynthetic apparatus (**Table 2**) in the S02oeFlv3 strain compared to the S02 strain, even at the initial lag phase of the growth on d1.5.

By d5.5, the S02oeFlv3 strain exhibited increased metabolic activity and cellular energy levels compared to the S02 control strain. This was evident from significantly enhanced growth (**Figure 2**) and sucrose production (**Figure 3**), as well as storage compound accumulation (**Figure 5**). The S02oeFlv3 strain also exhibited higher carbon uptake and oxygen evolution activity (**Figure 6**), as well as the further upregulation of sulfate transporters and the accumulation of several proteins involved in phosphate transport, ammonium uptake and assimilation (**Table 2**). These are all cellular processes with high ATP demand and an increased need for macronutrients. While we did not directly record increased ATP levels in the S02oeFlv3 strain, all these changes align with earlier observations that Flv3 overexpression in WT *Synechocystis* background leads to higher ATP availability, alongside enhanced O_2_ evolution and CBB cycle turnover (Hasunuma, et al., 2014). In the S02oeFlv3 strain, most of the carbon assimilated in CBB cycle is directed towards sucrose biosynthesis and other metabolic pathways, such as the oxidative TCA cycle, providing building blocks for growth. Storage compounds such as glycogen and PHB also accumulate already in d5.5 samples of S02oeFlv3 as compared to the S02 strain, but show further significant increase towards d9.5 when the cell growth begins to slow down. Interestingly, despite increased CO_2_ uptake and indications of nutrient imbalance, proteins typically associated with high-carbon sensing, such as the nitrate transporters involved in balancing high C/N ratio (Forchhammer and Selim, 2020) (Muth-Pawlak, et al., 2022) showed no difference between S02oeFlv3 and S02 (**Table 2**).

While the catalysis of ROS-independent O_2_ photoreduction is characteristic of *Synechocystis* Flv2/Flv4 and Flv1/Flv3 heterodimers, the data presented here suggest that Flv3 may also participate in an unknown redox reaction that does not use O_2_ as a substrate. Earlier studies have already shown that the four Flv paralogs have unique roles and interaction partner preferences in the typical active conformations, and that they cannot be interchanged without affecting the enzyme function. Our findings here, based on proteomic and biochemical data, corroborate the previously proposed idea that Flv3 can form catalytically active homomeric complexes that function differently to the Flv1/Flv3 counterpart. The distinct reactions between the Flv1/Flv3 heterodimer and the putative Flv3/Flv3 homodimer/oligomer therefore demonstrate that the Flv quaternary structure effectively determines the enzyme reaction specificity with regard to the preferred substrate. However, although the Flv3 homomer is expected to participate in an alternative photosynthetic electron transfer route from PSI, similar to the Flv1/Flv3 heterodimer, the final electron acceptor remains unknown. The acquired data on the sucrose-producing Flv3 overexpression strain indirectly suggests that this Flv3 homomer may be linked to sulfate metabolism, as discussed in more detail below.

### Sulfate assimilation as an electron sink from PSI in sucrose-producing Flv3 overexpression strain?

Sulfate assimilation is generally known to be coordinated with assimilation of carbon and nitrogen (Lappartient, et al., 1999) (Kopriva, et al., 2002) (Kopriva and Rennenberg, 2004). We observed an accumulation of sulfate transporters from d1.5 onwards, which increased significantly during the peak of growth and sucrose production at d5.5 in S02oeFlv3 **(Table 2)**. Compared to S02, the Flv3 overexpression strain appeared to adapt to increase sulfate assimilation from the outset, while the differences in the abundances of proteins and metabolites involved in carbon and nitrogen assimilation were minimal. These results suggest divergence in S02oeFlv3 sulfur metabolism, and changes in the coordination between the carbon, nitrogen and sulfate assimilation compared to S02.

The import and conversion of sulfate (SO₄²⁻) to sulfide (S₂⁻) (Karvansara, et al., 2024) requires significant cellular energy and reducing power. Sulfate is imported into the cell against the concentration gradient and requires activation by ATP to form adenosine 5’ phosphosulfate (APS). In the cytosol APS is converted to sulfide (S₂⁻) in the two reduction steps. The first reaction, two-electron reduction of APS to sulfite (SO₃²⁻) is catalyzed either directly by AP reductase (APR) or after APS phosphorylation to 3’phosphoadenosine 5’phosphosulfate (PAPS) by PAPS reductase (PAPSR). Reduced glutathione provides the necessary reducing power to both APR and PAPSR via thioredoxin or glutaredoxin. Consistent with this, we observed significant decrease in the levels of oxidized glutathione (d5.5) in S02oeFlv3 compared to S02 (**Table 3**), suggesting that the glutathione pool was predominantly reduced in this strain. The second reaction, six-electron reduction of SO₃²⁻ to S^2^⁻, is catalyzed by sulfite reductase (SiR), which uses ferredoxin (Fd) as the electron donor. Sulfide is then typically assimilated into O-acetyl-serine (OAS) to form cysteine (Takahashi, et al., 2011) which serves as a source of sulfur for subsequent metabolic reactions, and is the first metabolite to connect the assimilation pathways of the three major nutrients; carbon, nitrogen and sulfur. Sulfide is also an important redox element in the physiological network that allows for the modifications of proteins in the form of polysulfades (Shimizu, 2026) (Han, et al., 2022), as well as acting as a signaling compound in the form of H_2_S (Cuevasanta, et al., 2017). Interestingly, we observed the accumulation of SuoS light-dependent sulfide-quinone reductase (SQR) in the S02oeFlv3 strain of *Synechocystis* at d9.5. SQR is an enzyme that retrieves electrons from sulfide (i.e. hydrogen sulfide, H_2_S) and transfers them in light to the PQ pool in the thylakoid membrane (Nagy, et al., 2014) (**Figure 8**) simultaneously generating a polysulfide that then spontaneously reacts with glutathione (GSH) to form highly reactive glutathione persulfide (GSSR) and further glutathione disulfide (GSSG) (Wang, et al., 2024). The observed increase in oxidized glutathione content (d9.5) in S02oeFlv3 suggests the presence of reactive sulfur species (RSS), which, like ROS, are as harmful to cells. It seems likely that the S02oeFlv3 strain can use sulfate as an additional electron acceptor to support rapid cell growth and sucrose production. Overexpression of Flv3 in the sucrose-producing strain is associated with an increased photosynthetic rate and the need for an additional electron sink in the form of sulfates. Accumulation of reduced sulfur in the cell induces SQR accumulation, protecting the cell from toxic H_2_S and contributing to photosynthetic electron flow to PSI and respiratory pathways.

Harmful H_2_S may also be generated in the cell via a pathway involving the transformation of cysteine into pyruvate and persulfides (RSSH), which are subsequently transformed into polysulfides, releasing H_2_S via a series of enzymatic reactions (Han, et al., 2022) (**Figure 8**). In the S02oeFlv3 strain, accumulation of pyruvate and cyst(e)in desulfurase (SufS) (Kessler, 2004) indicates that this pathway is active in S02oeFlv3 and is strengthened by the accumulation of the SQR enzyme, which is necessary for the H_2_S detoxification. Interestingly, the SQR function in cyanobacteria is similar to the adaptation of certain bacteria to high H_2_S levels, allowing them to perform anoxygenic photosynthesis using H_2_S instead of H_2_O as an electron donor.

### Hypothetical function of Flv3 homooligomers in sucrose-producing *Synechocystis* cellular redox homeostasis

Based on our data, it is clear that the Flv3 homooligomer is formed and takes part in electron transfer reactions in *Synechocystis*. The ultimate metabolic electron and carbon sinks in sucrose-producing *Synechocystis* are biomass, sucrose, and the cellular storage compounds. While it is plausible that the electron donor for Flv3 homooligomer is ferredoxin, as in the case of Flv1/Flv3, the downstream electron acceptor from Flv3 homooligomer does not appear to be the molecular oxygen. Rather than wasting the electrons in the futile water-water cycle, as occurs with the Flv1/Flv3 heterodimer, it is suggested that the Flv3 homooligomer in the S02oeFlv3 strain effectively funnels the electrons for productive metabolic use (**Figure 8**). Based on all the collected proteomics and metabolomics data, sulfate reduction is the most promising candidate as a strong natural sink for reducing equivalents. Through this process, electrons can be directed, not only to metabolic processes such as cysteine biosynthesis, but also back to the LET (or CET) via reduction of PQ. Such a pathway is analogous to the anaerobic respiration performed by sulfate reducing bacteria that use SO_4_^2-^ as the terminal electron acceptor.

Taken together, we suggest that in *Synechocystis*, the Flv3 homooligomer serves as carrier between photosynthetic electron transport and sulfate reduction **(Figure 8)**. Unlike the Mehler-like reaction of the Flv1/Flv3 heterodimer, which channels the electrons to O_2_, the Flv3 homooligomer is suggested to relay the electrons for reduction of APS and SO_3_^2-^ via some yet unidentified redox interactions. When photosynthesis is active and there is an abundance of light, the electrons from the generated H_2_S can be rerouted to the PQ pool via light-regulated SQR. Thus, the sulfur metabolism provides an intermediate storage and a buffer for electrons derived from photosynthesis, which the cell can utilize for sulfur assimilation through O-acetyl-serine or for driving LET or CET through PSI via PQ or for respiration to generate ATP. This hypothesis is based on indirect phenotypic testing and omics data obtained from *Synechocystis* Flv3 overexpression strain engineered to produces sucrose as an efficient electron sink. Further biochemical experimentation is required to characterize putative molecular-level interactions.

## Author contributions

Conceptual planning by EMA, PK, DMP. Experimental design by EMA, PK, RN, DMP. Experimental work by RN, DMP, EM, AT. Experimental supervision by PK, RN, DMP. Data analysis and result interpretation by EMA, PK, RN, DMP. Manuscript preparation by EMA, PK, RN, DMP. Manuscript finalization by EMA, DMP. Preparation of Figures and Tables by RN, EMA, DMP, PK. Supervision and coordination by PK, EMA. All authors contributed to manuscript revision and approved the final version for publication.

## Acknowledgements

We performed the proteomic analyses at Turku Proteomics Facility, University of Turku and Åbo Akademi. The metabolomic measurements were made at FIMM Metabolomics Unit, Helsinki Institute of Life Science. Both facilities are supported by Biocenter Finland. The FIN-BioFoundry FIRI 2025-2028 national roadmap infrastructure, funded by the Research Council of Finland (decision # 367614), is acknowledged for providing research facilities and technical support.

## Supplementary Material

**Supplementary Figure S1:** Loading control for Figure 1 Flv3 immunoblot analysis (p. 2)

**Supplementary Figure S2:** Strain growth profiles when cultured in different volume (p.3)

**Supplementary Figure S3:** Calculated ratios between O_2_ photoreduction and PSII O_2_ evolution (p.4)

**Supplementary Table S1:** Summary of results from differential protein analysis expressed as log_2_FC values of S0oeFlv3 vs S02. (Excel file)

