## Supplementary Figures S1-S3 for "Overexpression of flavodiiron protein Flv3 in engineered *Synechocystis* stimulates sucrose production and growth by altering cellular redox balance through enhanced sulfur metabolism"

**Supplementary Figure S1:** Loading control for Figure 1 Flv3 immunoblot analysis (p. 2)

**Supplementary Figure S2:** Strain growth profiles when cultured in different volume. (p.3)

**Supplementary Figure S3:** Calculated ratios between $O_2$ photoreduction and PSII $O_2$ evolution (p.4)

**Supplementary Table S1:** Complete quantitative comparative proteomics data on S0oeFlv3 vs S02 (separate .xls)

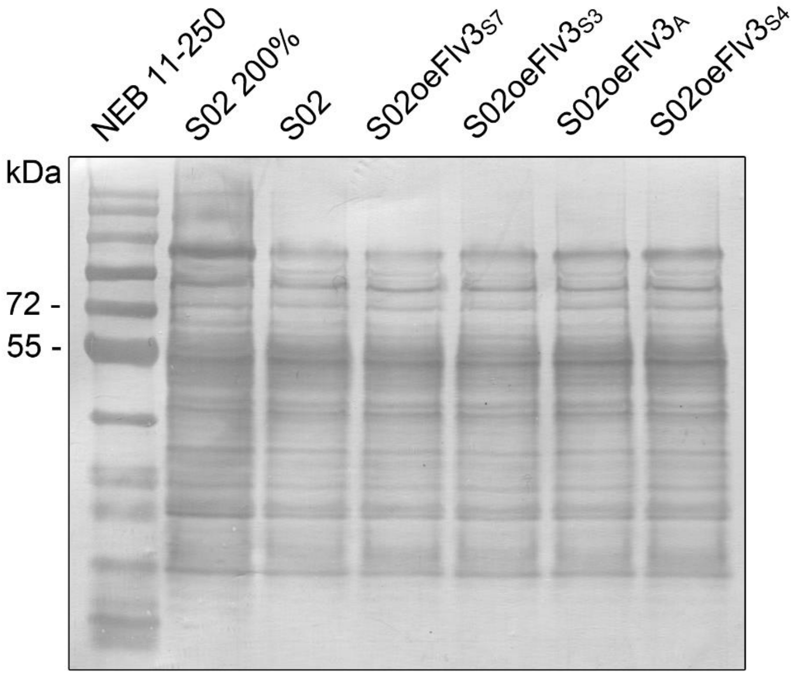


**Supplementary Figure S1: Loading control for Flv3 immunoblot analysis shown in Figure 1.** Coomassie-stained SDS-PAGE of the total protein extracts of the four S02oeFlv3 strains and the S02 control strain loaded at 15 µg total protein per well (100%) unless otherwise indicated. The calculated molecular weight of the Flv3 monomer is ~64 kDa. The ladder is NEB Blue Prestained Protein Standard (Broad Range 11-250 kDa).

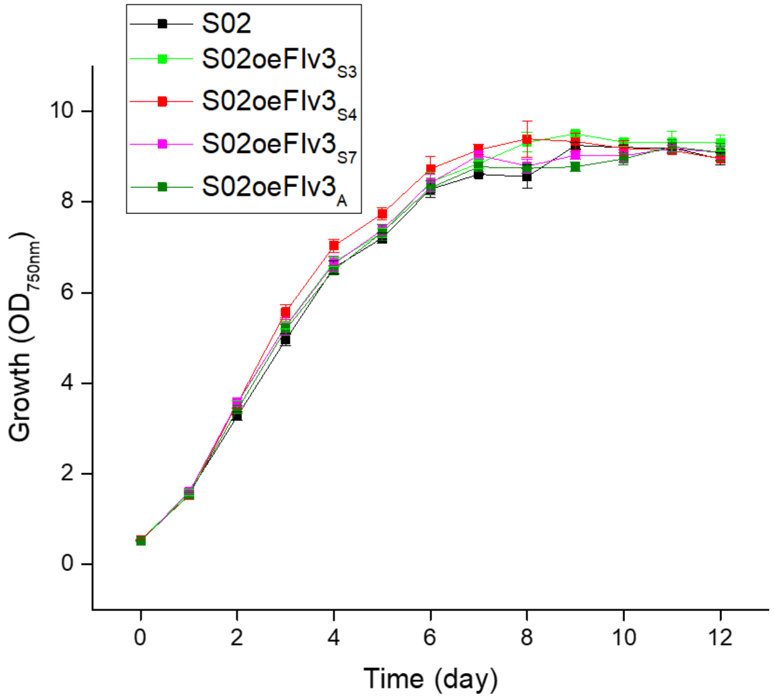


**Supplementary Figure S2:** Growth of the strains under 1% $CO_2$, 200 µmol photons $m^{-2}$ $s^{-1}$ continuous white light in BG11 in the presence of 400mM NaCl when cultured in 35ml medium volume in 100ml Erlenmeyer flasks. See Figure 2 for corresponding growth profiles in 100ml culture volume in 250ml flasks.

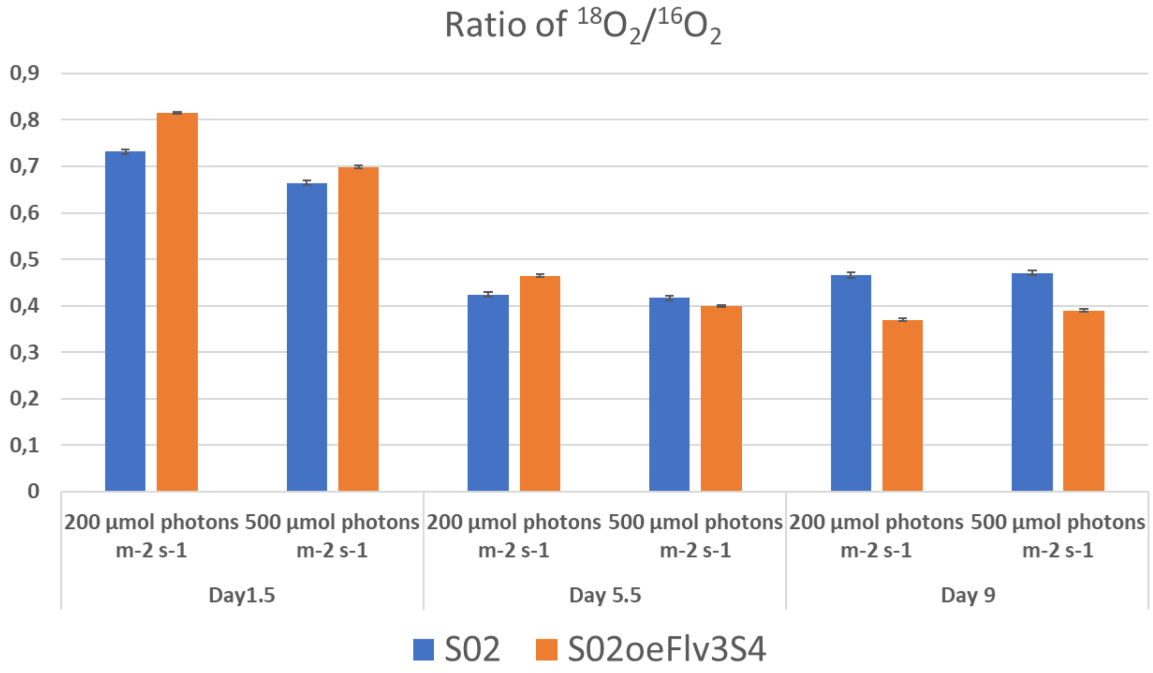


**Supplementary Figure S3:** The ratios between $^{18}O_2$ consumption ($O_2$ photoreduction) and $^{16}O_2$ evolution (the PSII oxygen evolution) calculated from the MIMS data (**Figure 6)** at three different sample points and at two light intensities. The bars represent the Flv3 overexpression strain S02oeFlv3$_{S4}$ (orange) and the S02 control strain (blue), calculated form three replicates.
